# Variation in neuronal activity state, axonal projection target, and position principally define the transcriptional identity of individual neocortical projection neurons

**DOI:** 10.1101/157149

**Authors:** Maxime Chevée, Johanna D. Robertson, Gabrielle H. Cannon, Solange P. Brown, Loyal A. Goff

## Abstract

Single-cell RNA sequencing technologies have generated the first catalogs of transcriptionally defined neuronal subtypes of the brain. However, the biologically informative cellular processes that contribute to neuronal subtype specification and transcriptional heterogeneity remain unclear. By comparing the gene expression profiles of single layer 6 corticothalamic neurons in somatosensory cortex, we show that transcriptional subtypes primarily reflect axonal projection pattern, laminar position within the cortex, and neuronal activity state. Pseudotemporal ordering of 1023 cellular responses to manipulation of sensory input demonstrates that changes in expression of activity-induced genes both reinforced cell-type identity and contributed to increased transcriptional heterogeneity within each cell type. This is due to cell-type specific biases in the choice of transcriptional states following manipulation of neuronal activity. These results reveal that axonal projection pattern, laminar position, and activity state define significant axes of variation that contribute both to the transcriptional identity of individual neurons and to the transcriptional heterogeneity within each neuronal subtype.

## INTRODUCTION

Single-cell RNA sequencing (scRNA-seq) approaches have been harnessed to reveal previously hidden levels of complexity in cell types and states within a given tissue^1, 2^. Nowhere is this more relevant than in the mammalian central nervous system, an organ system dependent on a remarkable diversity of cell types and cell states for its function. The neocortex contains a wide variety of neuronal cell types organized into the circuits that direct higher brain functions such as perception, memory and cognition. Furthermore, cortical neurons are highly dynamic, undergoing significant changes in gene expression during development and throughout adulthood in response to activity and experience^3-8^. Recent studies have generated a more comprehensive ‘parts list’ of discrete cell types within the neocortex than previously available^9-16^. While such surveys yield insights into the diversity of cortical cell types, the sources of transcriptional variation both within and across cell types remain poorly understood^17-19^.

We compared expression profiles of layer 6 corticothalamic neurons (L6CThNs), a heterogeneous population of related cortical projection neurons defined in anatomical, functional, and gene expression studies, making them ideally suited for investigating the relationships between transcriptional subtypes and other cellular properties^12, 16, 20-29^. By combining scRNA-seq with an enrichment strategy that preserved axonal target information, we identified two transcriptionally distinct L6CThN subtypes that each demonstrates a reproducible bias for long-range projection targets. The transcriptional profiles of L6CThNs also reflect their laminar position within L6. These two L6CThN subtypes exhibit divergent signatures of neuronal activity both in their baseline expression patterns as well as in their responses following manipulation of sensory input. The subtype biases in the choice of response following sensory manipulation increased transcriptional heterogeneity both within and between cell types and reinforced the transcriptional identities of the two L6CThN subtypes. These results demonstrate that scRNA-seq resolves relationships between gene expression and features such as axonal projection pattern, spatial organization, and cell state, and identifies the independent contributions of multiple biological signals that together determine transcriptional heterogeneity within and across neuronal populations.

## RESULTS

### Transcriptional profiling of layer 6 corticothalamic neurons reveals two subtypes which reflect axonal projection bias

Studies of primary sensory cortex demonstrate that L6CThNs are heterogeneous^20-29^. In rat barrel cortex, L6CThNs in upper layer 6 project to the ventral posterior medial nucleus (VPM) of the thalamus, while L6CThNs in lower layer 6 project to both VPM and the posterior medial nucleus (POm)^21, 25^. To distinguish between these two projection classes in mouse, we first validated Cre recombinase expression as a reliable marker for L6CThNs in the barrel cortex of *Neurotensin receptor 1-Cre* mice^30-32^ (Fig. 1a-e; *Ntsr1-Cre,* Gensat 220). Next, we showed that a subset of Cre-expressing, VPM-projecting L6CThNs in lower layer 6 also projects to POm (Fig. 1f-j).

**Figure 1:**
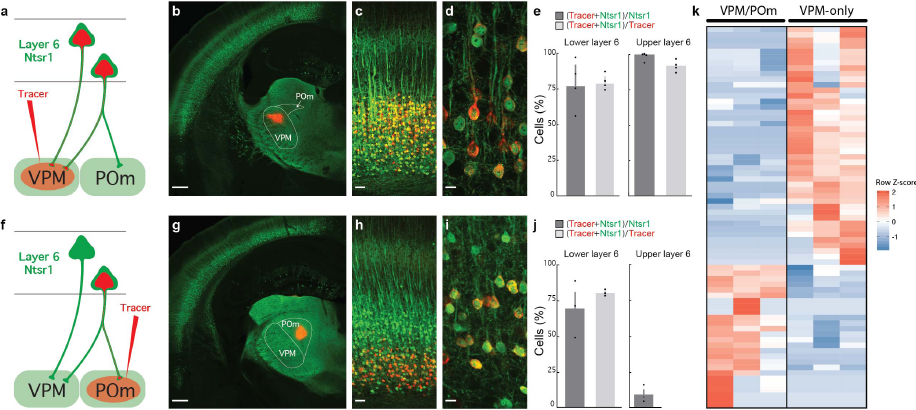
Layer 6 corticothalamic neuron subclasses distinguished by their axonal projections have distinct gene expression profiles. (**a**, **f**) The labeling schemes for two classes of layer 6 corticothalamic neurons (L6CThNs; VPM: Ventral posterior medial nucleus; POm: Posterior medial nucleus). (**b**) Injection of a retrograde tracer (red, Alexa 555-Cholera toxin B) into VPM in an Ntsr1-Cre;YFP mouse. Low (**c**) and high (**d**) magnification images of layer 6 (L6) of barrel cortex (BC) showing colocalization of the retrograde tracer (red) and YFP (green). (**e**) Quantification of the colocalization (mean±SE; n=4 mice). (**g**) Injection of a retrograde tracer (red) in POm of an Ntsr1-Cre;YFP mouse. Low (**h**) and high (**i**) magnification images of BC showing the colocalization of the tracer to CThNs in lower L6. (**j**) Quantification of the colocalization (mean±SE; n=3 mice). (**k**) Matrix showing the 69 genes differentially expressed between pools of VPM/POm and VPM-only L6CThNs from three replicates. Scale bars, (**b,g**) 500 μm; (**c,h**) 50 μm; and (**d,i**) 10 μm.

To determine whether L6CThNs subclasses projecting to VPM only or to both VPM and POm are distinguished by their gene expression profiles, we labeled the two subclasses in adolescent mice, microdissected barrel cortex containing retrogradely labeled cells, dissociated the tissue into a single-cell suspension, sorted the differentially labeled L6CThNs using Fluorescence Assisted Cell Sorting (FACS; Supplementary Fig. 1a-b), and collected enriched populations of each subclass for bulk RNA sequencing. We identified 69 genes with significant differential expression (DE) between the two sorted populations (Fig. 1k, Supplementary File 1, Supplementary Table 1; Cuffdiff2; 10% false discovery rate (FDR)). These 69 DE genes demonstrate that subclasses of L6CThNs distinguished by their long-range axonal projection patterns are differentiated by their gene expression profiles.

This bulk analysis was predicated on prior knowledge of existing morphological subclasses and may have obscured underlying transcriptional subtypes that comprise each projection class of L6CThNs. To resolve transcriptionally defined subtypes of L6CThNs, we next evaluated the gene expression landscape of single L6CThNs using an unbiased classification approach. We first sorted individual, labeled L6CThNs (Supplementary Fig. 1c) and collected 96 VPM-only and 96 VPM/POm L6CThNs from each of two replicate mice, totaling 384 single L6CThNs. Individual cell lysates were subjected to a modified Smart-Seq2 library preparation and scRNA-seq analysis. In total, 346 single L6CThNs passed quality control filters and were used for further analysis (Supplementary Fig. 1d-i; Supplementary File 2). We confirmed the fidelity of our enrichment by assessing each cell for neuronal and non-neuronal markers (Supplementary Fig. 1j).

To identify transcriptional subtypes of L6CThNs, we selected genes with the greatest likelihood of contributing to differences across the ensemble of single L6CThNs by identifying genes with high residuals to a mean-variance model fit independently in each of the two replicates and selecting the intersection of these two sets (Supplementary Fig. 2a). These 261 genes with high dispersion were therefore depleted for genes associated with technical variation between replicates^33, 34^ (Supplementary Table 2). Weights on the first three significant principal components across all cells using this gene set^35^ (Supplementary Fig. 2b; permutation parallel analysis; p < 0.001) were used for a tSNE dimensionality reduction followed by k-means clustering (Fig. 2a). Single-cell transcriptional profiles of the 346 L6CThNs clustered into at least two distinct subtypes. Fitting the data to three subtypes (k=3) or more did not significantly improve the performance of the clustering (Supplementary Fig. 2c). We compared our classification approach to several recently described single-cell clustering utilities^36, 37^ and found a high-degree of agreement (Supplementary Fig. 2d; SC3: 93.77% agreement; CIDR: 90% agreement). These results indicate that independent, unbiased clustering approaches based on genes with higher than expected variance across the L6CThN population identify two major subtypes of L6CThN.

**Figure 2:**
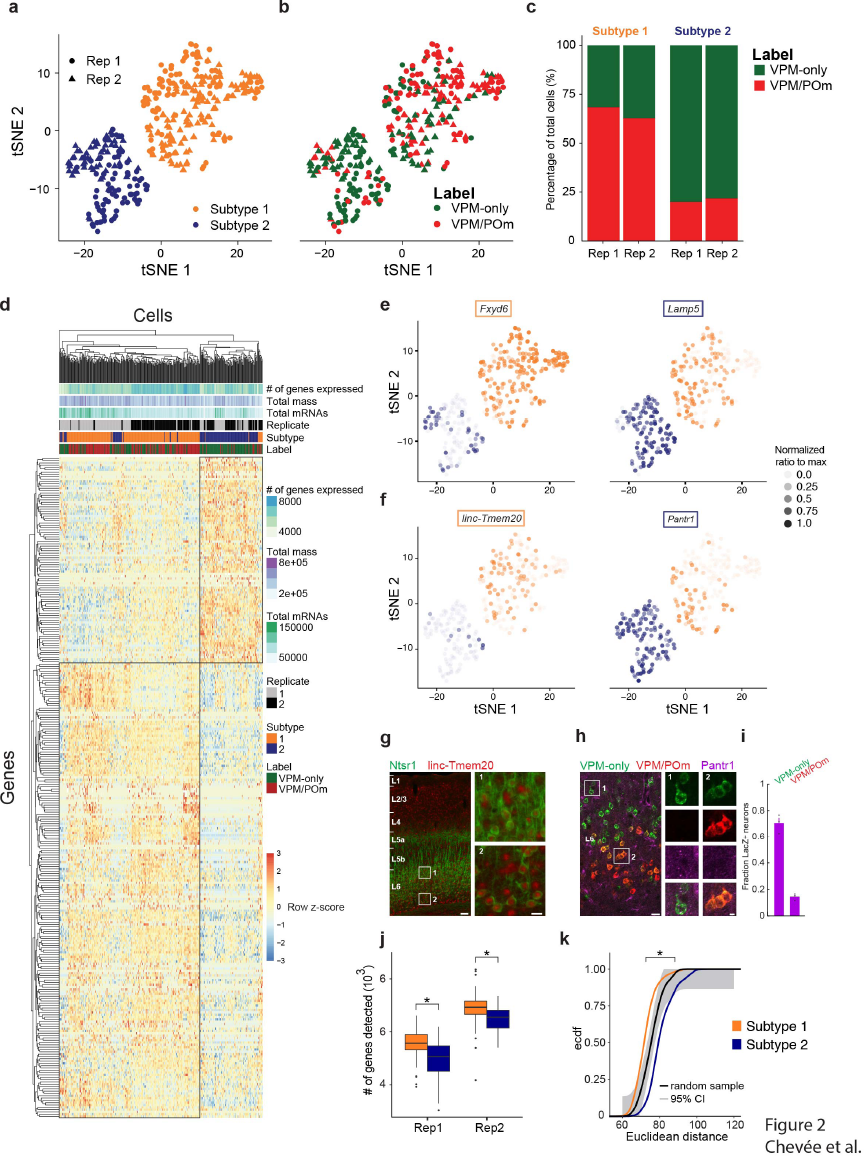
Unbiased clustering of single transcriptomes of layer 6 corticothalamic neurons defines two subtypes with strong axonal projection bias. (**a**) tSNE plot showing two subtypes of layer 6 corticothalamic neurons (L6CThNs) identified through unbiased clustering of single L6CThN transcriptomes. (**b**) Same tSNE plot as in (**a**), with each L6CThN color-coded by its axonal projection label. (**c**) Fraction of VPM-only (green) and VPM/POm (red) L6CThNs in each transcriptionally defined subtype for each replicate. (**d**) Matrix showing the hierarchical clustering of the 346 individual L6CThNs (x-axis) and the 286 genes differentially expressed (DE) between the two subtypes (y-axis, 0.1 % FDR). (**e**) tSNE plots showing the normalized expression levels of two differentially expressed genes enriched in subtype 1 *(Fxyd6,* left) or subtype 2 *(Lamp5,* right). (**f**) tSNE plots showing the normalized expression levels of two differentially expressed long-noncoding RNAs enriched in subtype 1 *(linc-Tmem20,* left) or subtype 2 *(Pantrl,* right). (**g**) Low magnification image of *linc-Tmem20* (red) in barrel cortex of an Ntsr1-Cre;tdTomato (green) mouse using a combination of *in situ* hybridization *(linc-Tmem20)* and immunohistochemistry (tdTomato). Insets show higher expression of *linc-Tmem20* in L6CThNs in lower layer 6 (L6, Inset 2) relative to upper L6 (Inset 1). (**h**) LacZ expression in barrel cortex of a heterozygous *Pantrl-LacZ* mouse following injections of green retrograde tracer in VPM and red tracer in POm. Insets show puncta of LacZ in VPM-only L6CThNs (Column 1, green) and not in VPM/POm L6CThNs (Column 2, red and green L6CThNs). (**i**) Barplot showing the fraction of VPM-only and VPM/POm L6CThNs expressing LacZ (mean±SE; n=3 mice). (**j**) Boxplot of the median number of genes detected across all cells for each subtype by replicate pair. (Replicate 1: Median number of genes detected: Subtype 1: 5582 ± 526.3 (sd), Subtype 2: 5080 ± 650.0 (sd), p < 2.169 × 10^−10^, Mann-Whitney test; Replicate 2: Median number of genes detected: Subtype 1: 6950 ± 545.4 (sd), Subtype 2: 6569 ± 478.7 (sd), p < 7.071 × 10^−7^, Mann-Whitney test). (**k**) Cumulative probability distribution of the pairwise Euclidean distances among cells in subtype 1 (gold) and subtype 2 (blue) (p < 2.2 × 10^−16^, Welch’s two-sample t-test). The black line represents the pairwise distances among a random sample of 100 cells drawn from the 346 cells. 95% confidence interval is shown in light grey (Dvoretzky-Kiefer-Wolfowitz inequality). Scale bars, (**g**) 100 μm, 20 μm; (**h**) 20 μm, 5 μm.

To determine the relationship between transcriptional identity and morphological subtypes, we next compared the distribution of VPM-only and VPM/POm projection labels across the two transcriptionally defined subtypes (Fig. 2b). The majority of neurons in subtype 2 were labeled VPM-only (79%, 103 of 130 cells, p < 3.768 × 10^−17^, hypergeometric test) while most neurons in subtype 1 were VPM/POm (65%, 141 of 216 cells, p < 3.768 × 10^−17^, hypergeometric test), a distribution significantly different from that expected by chance. Furthermore, the axonal projection bias was reproducible across replicates (Fig. 2c) even though the segregation of projection-defined subclasses was incomplete for both transcriptionally defined subtypes. Together our results indicate that each transcriptionally defined subtype of L6CThN is enriched for neurons targeting specific sets of thalamic nuclei.

### Transcriptional differences between subtypes of layer 6 corticothalamic neurons

To assess the transcriptional differences between the two identified L6CThN subtypes, all expressed genes were subjected to the Monocle2 differential test, using model formulas that accounted for both batch effects and the number of genes detected in each cell, a proxy measure for efficiency of RNA capture and library synthesis. We identified 286 genes that were significantly differentially expressed between the two subtypes (Fig. 2d; Monocle likelihood ratio test, 0.1% FDR; Supplementary File 2, Supplementary Table 3), only six of which overlapped with the 69 DE genes observed in our bulk RNA-seq analysis of sorted L6CThNs despite the high correlation between our bulk RNA-seq and scRNA-seq data (Supplementary Fig. 2e-g). This result, in conjunction with our previous observation of incomplete label segregation across L6CThN cell types, suggests that this discrepancy is primarily due to significant sample heterogeneity arising from retrograde label inefficiencies. Importantly, parameters such as total mapped fragments (mass), total estimated mRNAs per cell, number of genes detectably expressed per cell, and replicate did not result in biased clustering across the DE gene list (Fig. 2d), suggesting a minimal influence of technical variation on the list of DE genes. The significant differential expression of genes, including *Fxyd6* and *Lamp5,* between the two subtypes was consistent with expression patterns seen in the Allen Mouse Brain Atlas^38^ (Fig. 2e; http://mouse.brain-map.org, Fxyd6-RP_051017_01_E10-coronal, Lamp5-RP_050725_01_B03-coronal). The two subtypes also shared some transcriptional similarities with two recently defined subtypes in primary visual cortex^12^ (Supplementary Fig. 3). Several long non-coding RNAs (lncRNAs) were also specifically enriched in each subtype. For example, linc-Tmem20^8^ was significantly enriched in subtype 1 L6CThNs (Fig. 2f) and was preferentially expressed in lower layer 6 (Fig. 2g). Conversely, the lncRNA *Pantr1* was the gene with the greatest predictive power for neurons in subtype 2 (Fig. 2f, AUC = 0.876, power = 0.752, ROC analysis). Using a mouse line in which *LacZ* was knocked into the *Pantr1* locus^39^, we confirmed greater *LacZ* expression in VPM-only L6CThNs relative to VPM/POm L6CThNs (Fig. 2h,i). Furthermore, in contrast to our analysis of L6CThNs by anatomical labeling, we found that the total number of genes with detectable expression in each transcriptional subtype was significantly greater for subtype 1 relative to subtype 2, a finding consistent across replicates (Fig. 2j), and that the mean pairwise Euclidean distance between cells within each subtype was also greater in subtype 2 than in subtype 1, indicating greater cell-cell variation across subtype 2 (Fig. 2k; subtype 1 μ=72.16, σ=5.80; subtype 2 μ=80.59, σ=7.07; p < 2.2 × 10^−16^, Welch’s two-sample t-test). Importantly, we found only two genes, *Lypd1* and *Calm2,* with a significant combinatorial effect of subtype and label, suggesting that label does not distinguish subpopulations within each subtype. Together, these data identify two L6CThN subtypes and confirm a relationship between transcriptionally defined neuronal subtypes and the projection targets of these neurons.

To identify cellular processes that differentiate the two transcriptional subtypes of L6CThNs, we queried the DE gene list for enrichment of annotated gene sets from public databases (Supplementary Fig. 4a-d). Significant gene sets included Gene Ontology and Reactome terms related to general features of neurons such as “Neuronal part” and “Synaptic transmission” (p < 0.01; hypergeometric test, Benjamini-Hochberg corrected), highlighting the limited resolution of gene sets in currently available public databases for generating biological insights among neuronal subtypes. To identify more informative biological processes that shape differences in the response properties of L6CThNs, we compared the expression of voltage-gated ion channels, neurotransmitter receptors and neuropeptides, a number of which were differentially expressed (Supplementary Fig. 4e-h). Several G protein-coupled receptors and neuropeptides were differentially expressed. For example, *Adcyap1* (PACAP) and a gene encoding a peptide for processing PACAP, *Pam,* were preferentially expressed in subtype 2 (Supplementary Fig. 4h-i). Interestingly, receptors for PACAP are found in primary sensory thalamic nuclei^40^ and modulate thalamocortical interactions^41^, consistent with the projection pattern bias of subtype 2 L6CThNs. Our data indicate that a focused analysis identifies gene expression differences that reflect relevant functional features.

### Distinct cellular processes are coordinately regulated within subtypes of L6CThN

To identify cellular processes that contribute to the heterogeneity of gene expression across the transcriptomes of all L6CThNs analyzed, we performed a weighted gene co-expression network analysis (WGCNA) on all genes expressed in the 346 L6CThNs^42^. This analysis yielded 22 modules of co-regulated genes, which we classified using hierarchical clustering (Fig. 3a, Supplementary Fig. 5a). We tested for significant correlations between subtype identity for each cell and the eigenvalues of each module. Three modules significantly correlated with subtype 1 (Black, Turquoise, and Cyan; Fig. 3b, Supplementary Fig. 5b; p < 0.01; Pearson’s product moment correlation test) and four with subtype 2 (Red, Purple, Blue and Midnight Blue; Fig. 3b, Supplementary Fig. 5b; p < 0.01; Pearson’s product moment correlation test). When the same analysis was performed based on cell label, VPM-only or VPM/POm, correlation coefficients and confidence measures were all weakened (Fig. 3b, Supplementary Fig. 5c). These seven modules also showed significant correlation with the first principal component of the PCA on the high-variance gene set (Fig. 3c, Supplementary Fig. 3h), suggesting that subtype identity explains a significant amount of variation in gene expression across these neurons. Five of these modules were enriched for genes identified as significantly differentially expressed between subtype 1 and subtype 2 (Fig. 3d, Supplementary Fig. 3d). No module was correlated with replicate or other potentially confounding technical parameters (Supplementary Fig. 5f,g), confirming that the variations we observe are driven by biological differences among neurons rather than technical variation. Taken together, these data demonstrate that the greatest source of variation in the transcriptomes of L6CThNs is the difference between subtypes and reveal several discrete modules of gene expression contributing to this difference.

**Figure 3:**
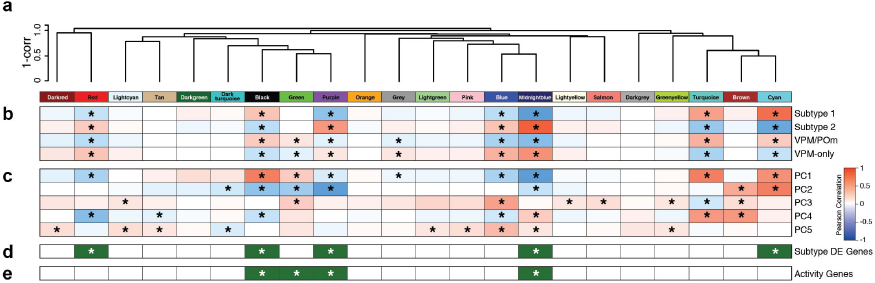
Coordinately regulated gene sets contribute to the transcriptional identities of layer 6 corticothalamic neuron subtypes. (**a**) Weighted gene co-expression network analysis (WGCNA) on variance-stabilized gene expression estimates identifies modules of coordinately regulated genes, which are grouped using hierarchical clustering of module eigengenes. (**b**) Pearson correlation of each module eigengene with parameters for both transcriptional subtype and label. Significance of each correlation was determined (*) using the Pearson’s product moment test (p < 0.01; Benjamini-Hochberg corrected). (**c**) Pearson correlation of each module eigengene with component rotations for principal components 1-5. (**d,e**) Hypergeometric test for enrichment of the 286 genes differentially expressed between transcriptionally defined L6CThN subtypes (**d**) and genes associated with neuronal activity (**e**) within each module (p < 0.01; Benjamini-Hochberg corrected).

### Projection-dependent and position-dependent gene expression differences contribute to the transcriptional identity of layer 6 neurons

Although the transcriptional signature of L6CThNs reflects projection target bias, axonal projection pattern and sublaminar position are confounded among L6CThNs. VPM/POm neurons are predominantly located in lower layer 6 while VPM-only neurons are biased towards upper layer 6 (Fig. 1h). Thus the gene expression differences we identified may represent differences in sublaminar position within layer 6 rather than axonal projection pattern *per se.* To test this hypothesis, we took advantage of the fact that L6CThNs represent only approximately half of the neurons in layer 6^29, 31^. If a gene’s expression reflects axonal projection pattern, its expression should be restricted to L6CThNs in upper or lower layer 6 only. On the other hand, if a gene’s expression reflects laminar position, we predict that its expression would be restricted to neurons in either upper or lower layer 6, regardless of their projection pattern.

To select target genes to evaluate using single molecule fluorescence *in situ* hybridization (smFISH), we performed a PCA on the mean-centered expression estimates of the high-variance genes across all 346 L6CThNs and used the rotations from this analysis to project all expressed genes into this PCA space to rank order candidate genes (Supplementary Fig. 6a-d). We quantified the expression of target gene mRNAs in individual tdTomato-positive, NeuN-positive neurons identified as L6CThNs and in tdTomato-negative, NeuN-positive neurons in slices of barrel cortex from Ntsr1-Cre;tdTomato mice. These data were then fit to a generalized additive model to test the independent contributions of laminar position and expression of tdTomato (Fig. 4). We found that *Lamp5, Serpini1* and *Gabra5* were selectively expressed in a subtype of L6CThNs (Fig. 4b,e-f). In contrast, *Pantr1* varied with laminar position within layer 6 in both L6CThNs and non-L6CThNs (Fig. 4c). Our findings reveal that information about position within layer 6 and long range axonal projection pattern is contained in the gene expression differences between the two transcriptional subtypes of L6CThNs.

**Figure 4:**
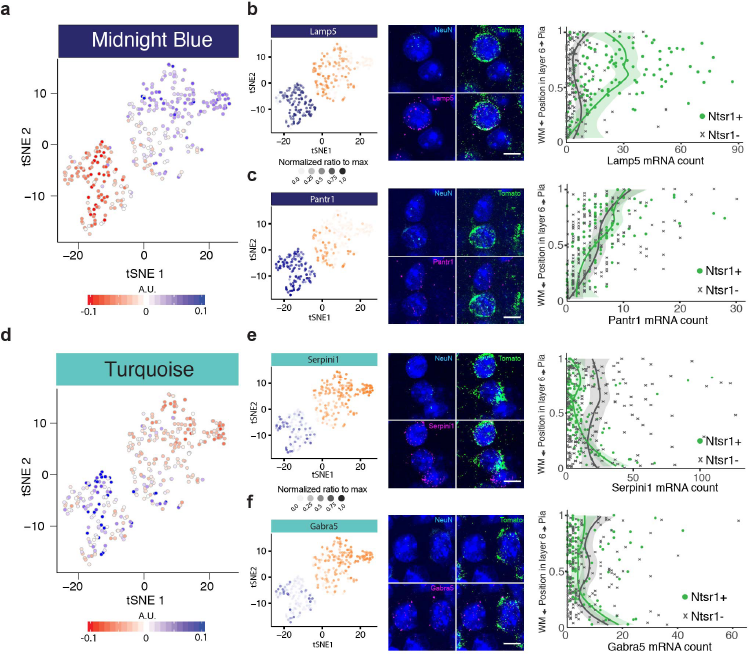
Variation in the transcriptional profiles of layer 6 corticothalamic neurons is defined by both subtype-specific genes and genes reflecting laminar location within layer 6. (**a**,**d**) tSNE plots showing the eigenvalue for each cell for the two WGCNA modules most correlated with the first principal component (**a**, Midnight Blue; **d**, Turquoise). (**b,c,e,f**) Gene expression tSNE plots *(left)* showing the normalized gene expression level in each cell for representative genes with significant weights on PC1. (**b**) and (**c**) are representative members of the Midnight Blue module, and (**e**) and (**f**) belong to the Turquoise module. Single molecule fluorescence *in situ* hybridization (smFISH, *middle)* showing images of mRNAs detected for each gene of interest (magenta), tdTomato (green), NeuN (cyan), and DAPI (blue) in layer 6 of Ntsr1-Cre;tdTomato mice. Quantitative gene expression analysis of smFISH *(right)* showing the number of mRNAs expressed per neuron as a function of normalized vertical position in layer 6 and neuronal cell type (L6CThNs: Ntsr1;tdTomato-positive;NeuN-positive neurons in green and non-L6CThNs: Ntsr1;tdTomato-negative, NeuN-positive neurons in grey). Curves represent Loess fits to the individual data points, grouped by cell type, and shaded areas correspond to 95% confidence intervals. Statistics: (**b**, *Lamp5)* “Subtype specific” p < 7.3231 × 10^−19^; “CThN+ position specific” p < 1.175 × 10^−17^; “CThN-position specific” p < 0.0035; (**c**, *Pantr1)* “Subtype specific” p < 0.9931; “CThN+ position specific” p < 9.243 × 10^−11^; “CThN-position specific” p < 1.021 × 10^−14^; (**e**, *Serpinil)* “Subtype specific” p < 1.606 × 10^−06^; “CThN+ position specific” p < 1.045 × 10^−06^; “CThN-position specific” p < 0.3342; (**f**, *Gabra5)* “Subtype specific” p < 0.1020; “CThN+ position specific” p < 1.994 × 10^−3^; “CThN-position specific” p < 0.09873.

### Neural activity significantly contributes to transcriptionally defined subtype identity

Neuronal activity is known to strongly influence gene expression^4-6^. We therefore hypothesized that neuronal activity state influences the transcriptional profiles of the L6CThNs and may contribute to their transcriptional identity. To test whether any modules reflect activity state, we assessed genes assigned to each module for enrichment of a curated gene set representing the ensemble of genes regulated after induction of neural activity compiled from recently published studies^43-46^ (Fig. 3e, Supplementary Fig. 5, Supplementary Table 4). Four of the seven modules correlated with transcriptional subtype (Black, Green, Purple, and Midnight Blue) were significantly enriched for genes induced by neuronal activity^43-46^ (Fig. 3e, Supplementary Fig. 5, p < 0.01, hypergeometric test). Each of these four modules was also significantly correlated with the first and second principal components of the PCA on the high-variance gene set, suggesting that differences in neuronal activity state explain a significant amount of transcriptional variation across L6CThNs as well. Taken together, our results show that the long range axonal projection pattern, laminar position within layer 6, and the activity state of each neuron are all reflected in the transcriptional profiles of individual L6CThNs and are principal contributors to the identity of the two L6CThN subtypes.

Among the four activity-associated modules (Black, Green, Purple, and Midnight Blue), Black was specifically correlated with subtype 1, and Purple and Midnight Blue with subtype 2, suggesting subtype-specific engagement of activity-induced genes in the steady state. The Green module was correlated with PC1-PC3, but demonstrated no significant bias for either cell type. To assess whether these signatures of neuronal activity represent a fundamental aspect of subtype identity, we re-evaluated our classification workflow after regressing out the eigenvalues for each activity-associated module and observed a reduction in the separation of the two L6CThN subtypes in each case, suggesting that steady state differences in neuronal activity genes are a defining characteristic of these two L6CThN subtypes (Supplementary Fig. 6e).

To further test the influence of neuronal activity on transcriptional identity, we evaluated the enrichment of activity genes along the first two principal components of a gene-centric PCA of the mean-centered expression estimates of high variance genes across all 346 L6CThNs. All expressed genes were projected in this PCA space and ranked using their weights along PC1 and PC2 (Supplementary Fig. 6a-d). Both PC1 and PC2 were significantly enriched for genes drawn from our curated list of activity-induced genes^43-46^ (p < 0.01; Kolmogorov-Smirnov test, Pre-ranked GSEA). The expression levels of the most heavily weighted PC1 genes varied predominantly between subtypes (Fig. 5b-c), while the expression levels of heavily weighted PC2 genes additionally varied within each subtype in a pattern similar to classical activity-induced genes like *Fos* and *Bdnf* (Fig. 5d-g), suggesting that activity contributes both to subtype identity and to transcriptional heterogeneity within each subtype. The expression patterns of genes such as *Cdh13, Igfbp6,* and *Sla* that strongly contributed to both PC1 and PC2 provide further evidence that axonal projection pattern, laminar location, and activity state are not fully orthogonal axes contributing to cellular identity (orange asterisks, Fig. 5c, Supplementary Fig. 6c,d). These data indicate that neuronal activity accounts for significant variation in the gene expression of L6CThNs, contributing to transcriptional heterogeneity both within and between L6CThN subtypes.

**Figure 5:**
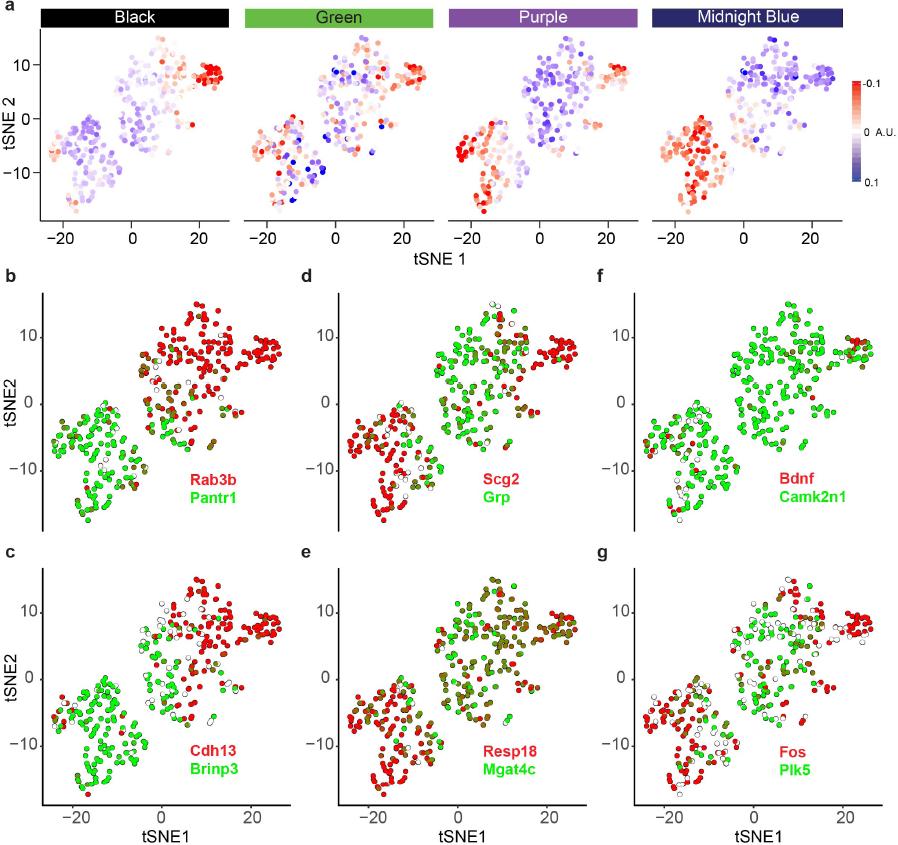
A signature of neuronal activity significantly contributes to the transcriptional identities of layer 6 corticothalamic neuron subtypes. (**a**) Module eigengenes for the four modules with significant enrichment for genes associated with neuronal activity. (**b,c**) tSNE plots showing the discretized expression of genes with the highest (**b**) and second highest (**c**) weights in each direction of PC1. (**d,e**) tSNE plots showing the discretized expression of genes with the highest (**d**) and second highest (**e**) weights in each direction of PC2. (f,g) tSNE plots showing the discretized expression levels of four genes reflecting neuronal activity: *Bdnf* and *Camk2n1* (**f**), and *Fos* and *Plk5* (**g**).

### Modulation of neuronal activity influences the transcriptional state and identity of L6CThNs

Our data demonstrate that the gene expression profiles of L6CThNs reflect an integration of multiple basis vectors corresponding to discrete, but potentially dependent sources of variation dominated by axonal projection pattern, sublaminar position within layer 6 and neuronal activity. Alterations in the molecular cascades engaged by different patterns of neural activity therefore have the potential to modulate the transcriptional identity of L6CThNs. To test this hypothesis, we unilaterally removed whiskers in a chessboard pattern, a pattern of sensory deprivation shown to engage plasticity mechanisms in the barrel cortex^47, 48^, from Ntsr1-Cre;tdTomato mice which were bilaterally injected with a retrograde tracer in POm to label VPM-only and VPM/POm L6CThNs as in our baseline experiments (Fig. 6a). At one and seven days following this sensory manipulation, we collected and sequenced single L6CThNs from the barrel cortex both contralateral and ipsilateral to the manipulation. After preprocessing and quality control, we obtained high-quality transcriptional profiles for 133 of 151 sequenced L6CThNs from 2 replicates one day following whisker removal (Day 1) and for 550 of 563 sequenced L6CThNs from 2 replicates seven days following whisker removal (Day 7). When combined with L6CThNs collected under baseline conditions (Day 0), we generated a data set comprised of 1023 sequenced L6CThNs collected from two replicates at each of three time points (Supplementary Fig. 7a-d, Supplementary File 3).

**Figure 6:**
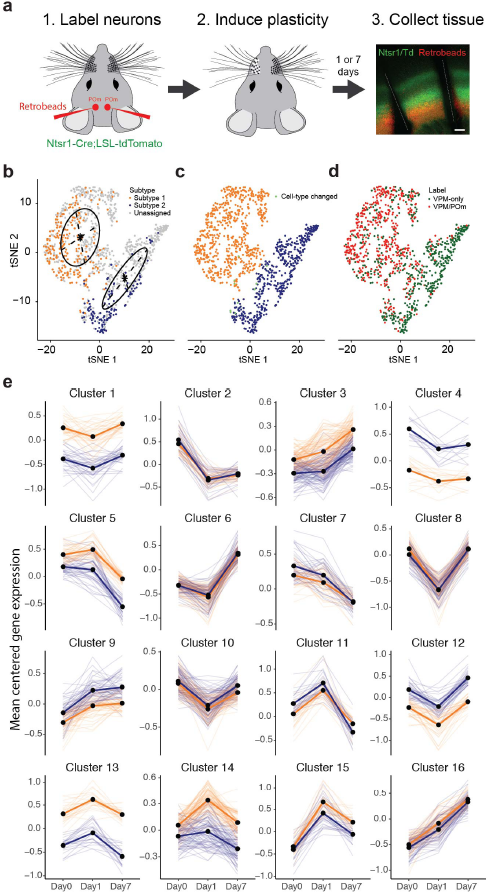
Sustained modulation of gene expression after sensory manipulation in layer 6 corticothalamic neurons. (**a**) Experimental design for sensory manipulation and single-cell transcriptional analysis of L6CThNs from the barrel cortex at 1 and 7 days post manipulation. (**b**) tSNE plot of all 1023 neurons obtained from baseline (Day 0), 1, and 7 days following sensory manipulation using the 286 genes identified as differentially expressed between L6CThN subtypes at day 0. Day 0 neurons are labeled by transcriptional subtype. Day 1 and Day 7 neurons are colored light gray. tSNE positions were fit to a Gaussian mixture model (black lines) which was used to classify Day 1 and Day 7 neurons as members of the previously defined L6CThN transcriptional subtypes. (**c**) tSNE plot colored by transcriptional subtype as assigned in (**b**). 10 of 340 neurons (2.9%) from Day 0 were assigned to a different subtype than in Fig. 2a and are labeled light green. (**d**) tSNE plot with neurons colored by projection label. (**e**) k-means clustering analysis of mean-centered gene expression, aggregated by Day and transcriptional subtype (Subtype 1: gold; Subtype 2: blue) for genes with significant differential expression after sensory manipulation. Semi-transparent lines represent individual genes and bold lines represent cluster centroids.

To assign each newly acquired L6CThN to its transcriptional subtype, gene expression profiles were first pre-processed and normalized as described for the baseline data set (Day 0). We then performed a PCA using the 286 genes we identified as differentially expressed between transcriptional L6CThN subtypes under baseline conditions and a tSNE analysis on the 1023 L6CThNs (Fig. 6b). Neurons were clustered using a Gaussian mixture model^49, 50^ (MClust), and cluster assignments were used to designate a subtype identity to each newly profiled neuron. This approach largely recapitulated our original subtype assignments, as only ten neurons classified under baseline conditions were assigned to a different subtype (Fig. 6c). The significant axonal projection bias of the two transcriptional subtypes was maintained as subtype 1 neurons were predominantly labeled VPM/POm (70.7%, 408 of 585 cells, p < 2.97 × 10^−71^, hypergeometric test) and subtype 2 VPM-only (84.2%, 369 of 438 cells, p < 2.97 x10^−71^, hypergeometric test), highlighting the reproducibility of our method and the robustness of our subtype assignment (Fig. 6d).

We identified 1134 genes with significant differential expression across the three time points sampled, independent of stable, baseline differences in expression between the two transcriptional subtypes (Fig. 6e, q ≤ 0.001; Monocle2 test). A k-means clustering analysis of mean expression profiles identified 16 clusters of genes with different temporal expression following deprivation, but failed to identify any clusters with significant subtype-specific effects over time (Fig. 6e). However, the transcriptional changes induced in response to altered sensory input are unlikely to be synchronous across all neurons collected at each time point. To describe the cellular responses to altered sensory input without the confounding effects of neurons in diverse states intermixed at each time point, we established a pseudotemporal ordering for the 1023 L6CThNs derived from the 1134 genes with significant differential expression across time points (Fig. 7a). Briefly, using the Monocle2 DDRTree algorithm, cells were arranged in an embedded graph representation in a reduced dimensional space. In this manner, cells with similar expression profiles across the 1134 target genes were positioned next to each other, and a traversal through the graph revealed the sequence of transcriptional changes that reflect progression through a given biological process. Through this projection, we captured a smooth representation of transcriptional modulation to altered sensory input. As expected, the distribution of L6CThNs along pseudotime generally followed the temporal order of collection following whisker removal (Fig. 7a,b). Nonetheless, neurons from each time point were found throughout pseudotime, confirming that the transcriptional response to sensory manipulation is asynchronous across the population of L6CThNs. Interestingly, we observed no bias in the distribution of L6CThNs across pseudotime when the neurons were grouped by hemisphere ipsi- or contralateral to whisker removal, suggesting that longer term transcriptional responses in L6CThNs from both hemispheres are similar (Supplementary Fig. 7e).

We identified 1507 genes that were differentially expressed across pseudotime at a much higher stringency than our aggregate analysis independent of transcriptional subtype designation (q ≤ 0.0000001; Monocle2), indicating that a significantly greater fraction of the transcriptome was identified as modulated by sensory manipulation using the increased resolution of pseudotemporal ordering. To identify cellular processes modulated along pseudotime, we clustered the normalized response curves of the differentially expressed genes (Fig. 7c, Supplementary Table 5) and queried each cluster for enrichment of annotated gene sets from public databases and our curated list of activity-induced genes. The cluster with the earliest changes in expression (Cluster 2) corresponded to significant downregulation of the activity-associated genes (p < 5.65 × 10^−7^, hypergeometric test) that were enriched in the four activity-associated modules contributing to cellular identity. In contrast, the cluster representing the late response (Cluster 3) corresponded to upregulation of genes associated with chromatin remodeling and reorganization as well as upregulation of lncRNAs, suggesting a slower epigenetic response to sensory manipulation. Interestingly, genes associated with long-term potentiation (LTP) from Cluster 2 *(Calm1, Calm2, Gria1, Gria2, Plcb1, Plcb4, Ppp1ca,* and *Ppp3r1)* were downregulated early in the response while LTP-associated genes in Cluster 3 *(Crebbp, Adcy1, Grin1, Prkcb, Ppp3ca,* and *Ppp3cb)* were upregulated towards the end of pseudotime, suggesting that non-overlapping subsets of genes in this single gene set are regulated at distinct phases of the L6CThN response to sensory manipulation. We also identified 75 transcription factors (TFs) with significant differential regulation after sensory manipulation including activity-associated TFs^5, 6^ such as *Arc, Fos, Ier2, Junb, Mef2c,* and *Nr4a1,* which were expressed early and downregulated over the course of pseudotime. Several TFs involved in neural development^3^ including *Neurod6, Fezf2, Mef2c,* and *Foxp2* were transiently expressed during the response, suggesting a regulatory relationship between activity-dependent plasticity and neural development.

**Figure 7:**
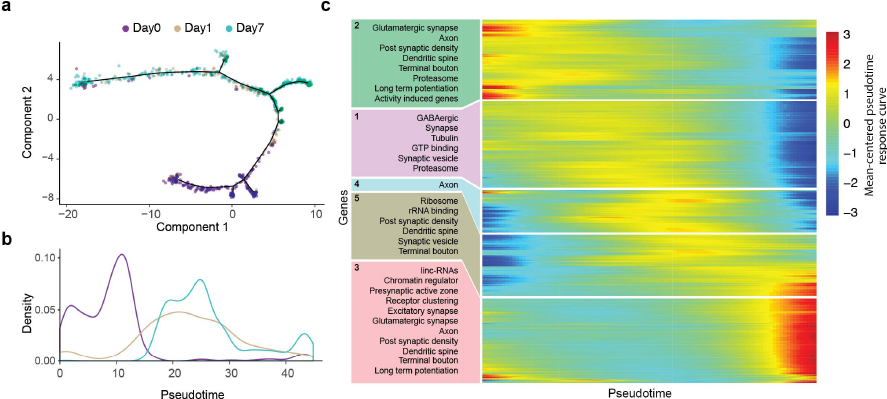
Pseudotemporal reconstruction of transcriptional responses to sensory manipulation in layer 6 corticothalamic neurons. (**a**) Discriminative dimensionality reduction projection of 1023 L6CThNs using genes identified as significantly differentially expressed after sensory manipulation. Neurons are colored by day relative to manipulation. (**b**) Density distribution of L6CThNs across pseudotime, grouped by day following manipulation. (**c**) Heatmap of normalized response curves for the 1507 genes with significant differential expression across pseudotime, and significantly enriched gene sets identified for each cluster (p < 1.0 × 10^−2^, hypergeometric test).

### Activity induced changes in gene expression enhance the distinction between subtypes

Our data demonstrate that a transcriptional signature of activity significantly contributes to L6CThN subtype identity under baseline conditions, and that altered sensory inputs result in dramatic transcriptional responses in L6CThNs. These results suggest that the response to sensory manipulation may alter the transcriptional relationship among neurons of the same subtype as well as the distinction between the two transcriptionally defined L6CThN subtypes. To test these hypotheses, we assessed the distribution of pairwise Euclidean distances of the variance stabilized gene expression estimates across all expressed genes for L6CThNs within each subtype for each day following sensory manipulation. In both subtypes, we observed that the response to altered sensory input resulted in an increased mean distance among cells across days (Fig. 8a; p < 2.2 × 10^−16^, Welch’s two-sample t-test). We also identified a significant increase in the variance of the intra-subtype distances across days (p < 3.59 × 10^−5^, F-test) for all adjacent time points except for subtype 2 Day 1 versus Day 7, indicating that the cell-to-cell variation within both L6CThN subtypes increases in response to altered sensory input. Second, we assessed the distribution of the inter-subtype pairwise Euclidean distances between subtype 1 and subtype 2. We found that the mean distance significantly increased from Day 0 to Day 1 as well as from Day 1 to Day 7 (Fig. 8b; p < 2.2 × 10^−16^, Welch’s two-sample t-test) as did the variance of inter-subtype distances (p < 4.67 × 10^−8^, F-test). These results demonstrate that modulation of neuronal activity increases both the transcriptional heterogeneity within each L6CThN subtype and the relative transcriptional differences between L6CThN subtypes, confirming a dependent relationship between activity level and transcriptional identity.

**Figure 8:**
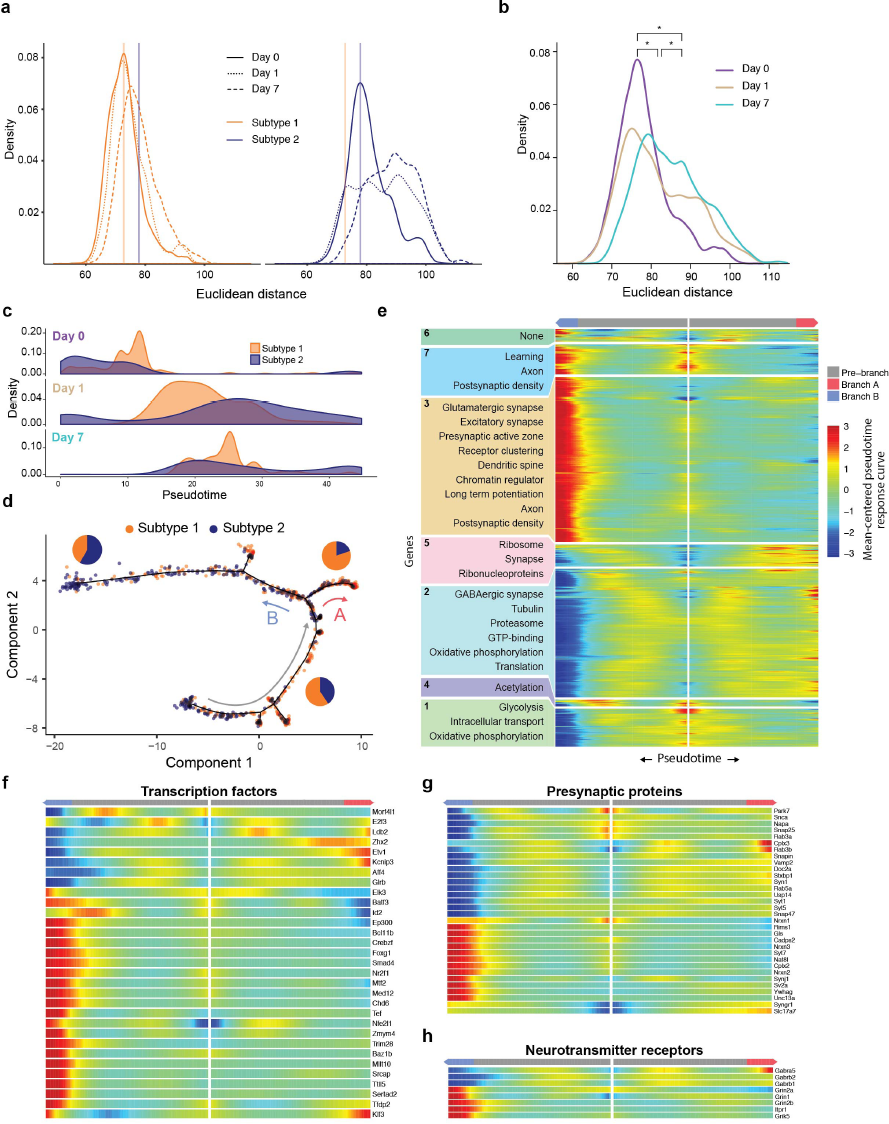
Sensory manipulation induces distinct cellular responses in layer 6 corticothalamic neurons biased with respect to transcriptional subtype. (**a**) Distribution of the pairwise Euclidean distances within each subtype for subtype 1 *(left,* gold) and subtype 2 *(right,* blue), using variance-stabilized expression estimates for all expressed genes, showing a significant increase in heterogeneity of gene expression for both subtype 1 and subtype 2. (**b**) Distribution of pairwise inter-subtype Euclidean distances between transcriptionally defined L6CThN subtypes across all expressed genes plotted for each day following sensory manipulation. There is a significant divergence between these two subtypes across time points as indicated by a positive shift in the distances after induction of experience-dependent plasticity. (**c**) Density distributions of L6CThNs at each day plotted across pseudotime for the two transcriptional subtypes of L6CThNs. (**d**) Discriminative dimensionality reduction projection of 1023 L6CThNs shown in Fig. 7a, now colored by transcriptional subtype. Red and blue arrows indicate the first major cellular response branch point following sensory manipulation. Grey arrow indicates the tree root state and direction of response progression. Pie charts indicate the proportion of each subtype in the population for each branch. (**e**) BEAM analyses of gene sets with significant differential expression dependent on either major branch point showing all significant DE genes. (**f-h**) BEAM heatmap for branch-dependent transcription factors not detected in the aggregate pseudotime response (**f**), presynaptic proteins (**g**), and ligand-gated neurotransmitter receptors (**h**).

### Subtype-biased responses contribute to transcriptional heterogeneity and subtype identity

These results raise the possibility that subtype-specific responses to sensory manipulation drive the increase in transcriptional heterogeneity and enhance the distinction between L6CThN subtypes. We found that subtype 1 and subtype 2 were differentially distributed along pseudotime at each time point assessed following sensory manipulation (Fig. 8c). Furthermore, pseudotemporal ordering identified a single major branch point in the transcriptional responses, with each of the two subsequent branches exhibiting a significant bias for a specific L6CThN subtype (Branch A and Branch B; Fig. 8d). Consistent with the proportion of the two subtypes in the baseline data, neither subtype 1 nor subtype 2 was significantly enriched in the root state (grey arrow; Subtype 1: p < 0.0715; Subtype 2: p < 0.91, hypergeometric test), indicating that both transcriptional subtypes share a similar early response to sensory manipulation. In contrast, Branch A and Branch B exhibited significant subtype bias: Branch A (red arrow) was biased for subtype 1 (VPM/POM; 80.4%, 148/184 neurons; p < 7.03 × 10^−14^, hypergeometric test) and Branch B (blue arrow) for subtype 2 (VPM-only; 58.4%, 202/346 neurons; p < 2.04 × 10^−13^, hypergeometric test). Similar subtype-specific biases were seen at a subsequent branch point (Supplementary Fig. 8). These results indicate that although either cell type may engage the discrete processes represented by each branch in response to sensory manipulation, the cellular decisions to engage a particular response are biased with respect to L6CThN subtypes.

Using the Monocle2 branch expression analysis modeling (BEAM) test, we identified 1392 genes with significant branch-dependent differential expression (Supplementary Table 6; q ≤ 0.0001; Monocle2 BEAM test); 926 of these overlapped with the 1507 genes with pseudotime-dependent expression suggesting that discrete cellular responses independently contribute to the aggregate transcriptional response to altered sensory manipulation. The branch-dependent genes were organized into seven distinct clusters using hierarchical clustering of the Monocle2 branched model fits (Fig. 8e). Hypergeometric testing demonstrated that neurons that progress along Branch A (VPM/POM enriched) were enriched for genes associated with engagement of the proteasome complex, a process involved in synaptic remodeling^51^, while Branch B cells (VPM-only enriched) were significantly enriched for genes related to the organization of the postsynaptic density as well as to the regulation of LTP. We identified 79 TFs expressed in a branch-dependent manner, including 31 TFs not identified in our initial analysis of the aggregate response (Fig. 8f), suggesting that much of the aggregate L6CThN response induced by altered sensory inputs is confounded across these two alternative cellular responses. The remaining 27 pseudotime-dependent TFs may regulate a uniform response independent of these two subtype-biased responses. Additional genes regulated in a branch-specific manner included thirty-two presynaptic genes (Fig. 8g). Furthermore, neurons committing to a Branch A response demonstrated enhanced expression of GABA receptor subunits *(Gabra5, Gabrb1* and *Gabrb2)* while neurons along Branch B induced ionotropic glutamate receptors (Fig. 8h; *Grin1, Grin2b, Grik5,* and *Grin2a).* Taken together, these subtype-specific biases in transcriptional responses induced by sensory manipulation likely underlie the overall effect we observed on L6CThN identity, enhancing the distinctions between subtype 1 and subtype 2, and significantly increasing both the inter-and intra-subtype heterogeneity of L6CThNs.

### DISCUSSION

Using scRNA-seq, we identified two transcriptionally defined subtypes of L6CThNs, each exhibiting significant bias for particular long-range axonal projection targets. Analysis of gene expression differences among the profiled neurons determined that variation in axonal projection pattern, laminar position within the cortex, and neuronal activity state all significantly contribute to transcriptional identity, and that these properties are at least partially dependent. Manipulating the activity states of L6CThNs by altering tactile input further showed that each subtype was biased for particular cellular responses to sensory manipulation, revealing functional differences among these neurons. These subtype-biased transcriptional responses not only increased cell-to-cell transcriptional heterogeneity within each subtype, but also enhanced the transcriptional differences between the two subtypes. Together, these data identify the most significant influences on the transcriptional identity of individual cortical projection neurons, and suggest that discrete cellular properties and responses affect the population-level variation and identity of neuronal subtypes.

Although it is possible that projection target and transcriptional identity are decoupled for a minority of neurons, the incomplete segregation of retrograde label across subtypes observed here likely represents mislabeling of a subset of L6CThNs, as expected due to the close proximity of VPM and POm and the difficulty of retrogradely labeling all neurons projecting to adjacent targets. Interestingly, the subtypes identified here share some transcriptional similarities with two recently defined subtypes in primary visual cortex^12^ (Supplementary Fig. 3), suggesting a conservation of corticothalamic neuron transcriptional identity and axonal projection target bias across functional regions of the cortex^20^. Future studies will be required to determine whether VPM-only and LGN-only projecting neurons as well as VPM/POM and LGN/LP neurons are transcriptionally analogous, and to identify sources of cellular variation across cortical areas. Although the population of L6CThNs we analyzed is comprised of two subtypes, profiling a greater number of neurons or profiling at greater depth may reveal additional rare subtypes of L6CThNs or may show that subtle variations in axonal projection pattern identified in some anatomical studies are not apparent in the expression profiles of cortical neurons^20, 21^.

Pseudotemporal ordering of the states induced in L6CThN transcriptomes by altered sensory input assessed over seven days revealed that L6CThNs engage at least two molecularly distinct responses in a subtype-biased, but not subtype-specific, manner. Although the distinct transcriptional responses were dominated by neurons collected one and seven days following whisker manipulation, neurons collected in the baseline state were found throughout pseudotime, suggesting that individual cortical neurons may engage these plasticity responses in the non-manipulated state. L6CThNs in the initial phase of cellular response to altered sensory input exhibited similar gene expression changes, indicating that both subtypes engaged a common initial transcriptional response. These results are consistent with previous studies of responses induced by neural activity measured over hours across brain regions and between inhibitory and excitatory neurons which found that common early transcriptional responses lead to cell-type specific late responses^52, 53^.

The differential response to sensory manipulation resulted in an increase in the transcriptional variation across L6CThNs within each subtype as well as a significant enhancement of the distinction between the subtypes, demonstrating a non-dissociable relationship between neuronal identity and neuronal activity. Because single L6CThNs have the potential to engage either transcriptional response regardless of subtype, our data suggest that extrinsic factors such as distinct activation patterns, due to differences in the circuits in which each subtype is embedded, induce neurons from a given subtype preferentially toward a similar response rather than solely cell-autonomous properties. Interestingly, we found that the transcriptional responses for neurons ipsilateral and contralateral to the sensory manipulation were similar, suggesting that shared cellular mechanisms underlie the previously described functional effects ipsilateral and contralateral to unilateral manipulation at timescales similar to those assessed here^48, 54, 55^. Furthermore, expression of genes that strongly contributed to subtype identity such as *Lamp5* was altered in response to sensory manipulation. These data indicate that factors that change cell state such as plasticity or injury affect our ability to accurately define discrete, stable transcriptional subtypes.

Our results indicate that the transcriptional profiles of cortical neurons reflect specific features of these cells, but also that the transcriptional variation across individual neurons is a principal feature of subtype identity with significant functional consequences for individual neuronal responses and subtype function. Subtype 2 L6CThNs were more transcriptionally heterogeneous than subtype 1 neurons under steady-state conditions in part because of baseline differences in gene expression associated with neural activity. In addition, the intra-cell type variation across subtype 2 at day 1 and at day 7 was greater than the inter-cell type variation at the same time points, suggesting that changes in cell-to-cell variation, rather than subtype specific differences, dominate the transcriptional responses to experience dependent plasticity. Together, these data indicate that the contribution of neuronal activity to gene expression differs across distinct neuronal subtypes and that the transcriptional variation due to differences in neuronal activity state plays a central role in defining the identity of cortical projection neurons.

## METHODS

### Mice

All procedures were approved by the Johns Hopkins Animal Care and Use Committee and followed the guidelines of the Society for Neuroscience and the National Institutes of Health. Animals used for RNA-sequencing ranged from postnatal day 23 (P23) to P28; animals used for immunohistochemistry and *in situ* hybridization ranged from P23 to P32. Both males and females were used in this study (gender indicated for each replicate below). All animals were kept on a 12 h light/dark cycle, housed with at least two mice per cage and provided with unlimited water and food. The following mouse lines were used:

Neurotensin receptor-1 Cre recombinase line^30^ (Ntsr1-Cre, GENSAT 220), loxP-STOP-loxP-tdTomato Cre reporter lines^56^ (Ai9 and Ai14, Allen Institute for Brain Science), loxP-STOP-loxP-eYFP Cre reporter line^56^ (Ai3, Allen Institute for Brain Science), and Pantr1-LacZ^39^. Mice were maintained on mixed backgrounds including C57BL/6 and CD-1.

### Stereotactic injections

To identify layer 6 corticothalamic neurons (L6CThNs), mice P18 to P23 were anesthetized with ketamine (50 mg/kg), dexmedetomidine (25 μg/kg) and the inhalation anesthetic, isoflurane. Animals were placed in a custom-built stereotactic frame and anesthesia was maintained with isoflurane (1-2.5%). A small craniotomy was performed, and a glass pipette (10-25 μm tip diameter) was lowered into the thalamus. To identify L6CThNs that project to the ventral posterior medial nucleus (VPM) of the thalamus, Ntsr1-Cre;tdTomato mice were injected in VPM (1.1 mm posterior, 1.7 mm lateral, 1.4 mm ventral from bregma) with either Alexa 488 Cholera toxin B (CTB488, ThermoFisher) or green Retrobeads IX (Lumafluor), and Ntsr1-Cre;YFP mice were injected with Alexa 555 Cholera toxin B (CTB555, ThermoFisher). To identify L6CThNs that project to both VPM and to the posterior medial nucleus of the thalamus (POm), Ntsr1-Cre;tdTomato mice were injected in POm (1.35 mm posterior, 1.23 mm lateral, 3.3 mm ventral from bregma) with green Retrobeads IX, and Ntsr1-Cre;YFP mice were injected with either CTB555 or red Retrobeads (Lumafluor). Between 30 and 100 nl of tracer were pressure-injected, and the pipette was kept in position for 5-10 minutes before removal. Following the injection, the incision was sutured and buprenorphine (0.05 mg/kg) was administered to all animals postoperatively. Quantification of the colocalization between retrogradely labeled neurons and Cre expression in Ntsr1-Cre mice (Fig. 1e,j) was performed on mice injected with either CTB or Retrobeads. Preliminary experiments indicated that all tracers used generated similar labeling. However, Retrobeads resulted in increased discrimination of labeled neurons during Fluorescence Assisted Cell Sorting (FACS) and were therefore used for all FACS experiments.

### Immunohistochemistry

Brains from animals injected with neuronal tracers were removed and placed in a solution of 4% paraformaldehyde (PFA) in 0.01 M phosphate buffered saline (PBS) for 3 h. Coronal sections were cut on a vibratome (50 μm, VT-1000S, Leica). Mice expressing YFP were subjected to immunohistochemistry (1:1000, chicken anti-GFP, GFP-1020, Aves, RRID:AB_10000240, and 1:300 AlexaFluor 488-conjugated goat anti-chicken, Life Technologies, A11039, RRID:AB_10563770). Sections were then mounted using Aqua Poly/Mount mounting medium (Polysciences, Inc) and visualized on a confocal microscope (LSM 510, Zeiss) using 10x (0.3 NA), 25x (0.8 NA) or 40x (1.3 NA) objectives or on an epifluorescence microscope (AxioObserver.Z1, Zeiss) using a 5x (0.16 NA) objective. Colocalization of tdTomato, YFP and retrograde tracer was quantified using single-plane confocal images and the Cell Counter plugin in Fiji^57^. For all anatomical comparisons, upper layer 6 and lower layer 6 were defined as the top 50% and bottom 50% of the layer defined by Cre-expressing neurons in Ntsr1-Cre mice.

### Cell isolation and enrichment

For bulk sequencing experiments (bulk RNA-seq), POm was injected with green Retrobeads in three Ntsr1-cre;tdTomato mice. For single-cell sequencing experiments (scRNA-seq), POm was injected with red or green Retrobeads in Ntsr1-Cre;YFP mice or in Ntsr1-Cre;tdTomato mice, respectively. Mice were sacrificed 5-7 days after surgery (Bulk RNA-seq: Replicate 1: P23 female; Replicate 2: P23 female; Replicate 3: P26 male; scRNA-seq baseline: Replicate 1: P27 male, Replicate 2: P28 female; scRNA-seq Day 1: Replicate 1: P24 female, Replicate 2: P24 female; scRNA-seq Day 7: Replicate 1: P29 female, Replicate 2: P27 female). Brains were rapidly removed and 300 μm thick somatosensory thalamocortical^58^ or coronal slices were sectioned (VT-1200s, Leica) in ice-cold sucrose solution containing the following (in mM): 76 NaCl, 25 NaHCO_3_, 25 glucose, 75 sucrose, 2.5 KCl, 1.25 NaH_2_PO_4_, 0.5 CaCl_2_, 7 MgSO_4_, pH 7.3, 310 mOsm continuously bubbled with 95% O_2_/5% CO_2_. The slices were then placed in a submersion chamber on an upright microscope in ice-cold sucrose solution continuously bubbled with 95% O_2_/5% CO_2_. The barrel cortex was identified using differential interference contrast (DIC) and epifluorescence microscopy (Axioskop2 FsPLus, Zeiss) using 4x (0.1 NA) and 40x (water immersion, 0.75 NA) objectives. The location of the retrograde tracer in the lower half of layer 6 was visually confirmed, and then regions of cortex below barrels with appropriate labeling were microdissected and immediately dropped into 2.6 mL equilibrated Papain DNase-I solution. For all experiments, at least three consecutive slices confirmed as clearly labeled were microdissected.

The dissociation protocol was adapted from the trehalose-enhanced neuronal isolation protocol described in Saxena et al, 2012^59^ which is based on the Worthington Papain Dissociation System (Worthington Biochemical Corporation). Briefly, tissue pieces were dropped into a 5 ml tube containing 2.6 mL of a trehalose-Papain-DNase1 solution which included RNase inhibitor, incubated 30-45 min at 37°C with gentle trituration at the half way point using a p1000 pipetter. The reaction was stopped by adding a protease inhibitor solution (for solution compositions, see Saxena et al, 2012^59^. The tissue was then further dissociated and washed by sequential pipetting and low-speed centrifugations. The final pellet was resuspended in 200 μL of media (DMEM, 5% trehalose (w/v), 25 μM AP-V, 0.4 mM kynurenic acid, 6 μL of 40 U/μl RNase inhibitor) at room temperature and introduced into a FACS machine (Beckman Coulter MoFlo Cell Sorter). Neurons were sorted based on their fluorescence (either tdTomato-positive (red) and Retrobead-positive (green) versus tdTomato-positive only or YFP-positive (green) and Retrobead-positive (red) versus YFP-positive only) directly into lysis buffer either in eppendorf tubes for bulk sequencing (350 μL lysis buffer, Qiagen RNeasy Micro Kit; Replicate 1: tdTomato-only: 1943 cells, tdTomato and tracer: 2287 cells; Replicate 2: tdTomato-only: 386, tdTomato and tracer: 782; Replicate 3: tdTomato-only: 604, tdTomato and tracer: 934) or into individual wells of 96-well plates for single-cell sequencing (2 μL Smart-Seq2 lysis buffer + RNAase inhibitor, 1 μL oligo-dT primer, and 1 μL dNTPs according to Picelli et al, 2014^60^. Upon completion of a sort, the plates were briefly spun in a tabletop microcentrifuge and immediately placed on dry ice. Single-cell lysates were subsequently kept at-80°C until cDNA conversion.

### Library preparation and amplification

For bulk samples, RNA was extracted with a Qiagen RNeasy Micro Kit. RNA quality was assessed with the Bioanalyzer Pico RNA kit and samples had RIN scores of 8.2-8.8. Library preparation and amplification were performed according to the Smart-Seq2 protocol^60^ with some modifications. Principally, all primer and adapter sequences were synthesized with a 5’-biotin to minimize primer dimerization. Briefly, 200 pg of RNA per sample were used as input for template switching cDNA synthesis. Full-length cDNA was amplified by KAPA HiFi mediated PCR for 20 cycles using a 5’-biotinylated ISPCR primer. Then, 500 pg of Ampure XP bead cleaned (1:1) cDNA was used as input for a standard Nextera XT tagmentation reaction, and amplification of adapter-ligated fragments was carried out for 12 cycles during which individual index sequences were added to each distinct sample. Library concentration was assessed with Qubit and library fragment size distribution was assessed on the Agilent Bioanalyzer. Pooled, indexed bulk RNA-seq samples were initially sequenced on the Illumina MiSeq 500 platform, and subsequently the same pool was sequenced on one lane of the Illumina HiSeq 2500 platform to produce 50 bp paired-end reads. Libraries were sequenced to an average depth of 4.2x10^7^ fragments each. Reads from both runs were aggregated by index prior to mapping.

Library preparation and amplification of single-cell samples was performed using a modified version of the Smart-Seq2 protocol. Briefly, 96-well plates of single-cell lysates were thawed to 4°C, heated to 72°C for 3 minutes, and then immediately placed on ice. Template switching first-strand cDNA synthesis was performed as described above using a 5’-biotinylated TSO oligo. cDNAs were amplified using 18 cycles of KAPA HiFi PCR and 5’-biotinylated ISPCR primer. Amplified cDNA was cleaned with a 1:1 ratio of Ampure XP beads and 300 pg was used for a one-half standard-sized Nextera XT tagmentation reaction. Tagmented fragments were amplified for 12 cycles and dual indexes were added to each well to uniquely label each library. Concentrations were assessed with Quant-iT PicoGreen dsDNA Reagent (Invitrogen) and samples were diluted to ~2 nM and pooled. Pooled libraries were sequenced on the Illumina HiSeq 2500 platform to a target mean depth of ~8.0 × 10^5^ 50 bp paired end fragments per cell^61^ at the Hopkins Genetics Research Core Facility to generate 50 bp paired-end reads.

### In situ hybridization and immunohistochemistry

Immunohistochemistry (IHC) for tdTomato and *in situ* hybridization (ISH) for *linc-Tmem20* were combined to show co-localization of the lncRNA transcript and tdTomato protein in L6CThNs. Ntsr1-Cre;tdTomato mice (P23 to P30) were transcardially perfused with 0.1 M PBS followed by a 4% PFA solution, and brains were post-fixed for 3 h at room temperature in 4% PFA. Then, 30 μm sections were cut on a vibratome (VT-1000S, Leica) and first processed for IHC (1:1000, Rabbit anti-DsRed polyclonal antibody, Living Colors, Clontech Laboratories, RRID: AB_10013483 and 1:300 AlexaFluor 568-conjugated goat anti-rabbit, Life Technologies, A11011, RRID: AB_143157). Next, the sections were processed for ISH following the protocol described in Blackshaw, 2013^62^ without proteinase K treatment. Sections were incubated overnight at 65°C with 5 ng/μL of an anti-sense *linc-Tmem20* DIG labeled probe (DIG RNA labeling Kit (SP6/T7), Roche). Detection of the hybridized DIG probes was performed using an alkaline phosphatase (AP) conjugated antibody (1:1000, Anti-Digoxigenin [21H8] Alkaline Phosphatase, Abcam, Cat. No.: ab119345, RRID:AB_10901703) followed by application of the AP substrate BCIP/NBT (SigmaFast BCIP/NBT, Sigma-Aldrich). The enzymatic reaction was carried out until the desired signal to noise ratio was observed. Images were then taken on an AxioObserver.Z1 (Zeiss) using a combination of epifluorescence and brightfield imaging (10x objective, NA: 0.3). Brightness and contrast were adjusted using Adobe Photoshop (Adobe Systems Incorporated).

For validation of *Pantr1* expression, transgenic mice expressing LacZ driven by the endogenous *Pantr1* promoter locus^39, 63^ were injected with CTB488 in VPM and CTB555 in POm. IHC was performed on 50 μm-thick sections (rabbit anti-β galactosidase antibody 1:5000; a kind gift from Dr. Joshua R. Sanes, Harvard University, Cambridge, MA and 1:300 AlexaFluor 647-conjugated donkey anti-rabbit, Life Technologies, A31573, RRID: AB_2536183). Imaging was performed on a confocal microscope (LSM 510, Zeiss) using 10x (0.3 NA), 25x (0.8 NA) or 40x (1.3 NA) objectives. LacZ-positive and retrogradely labeled neurons were manually counted on single-plane confocal images using the Cell Counter plugin in Fiji^57^. Brightness and contrast were adjusted using Adobe Photoshop (Adobe Systems Incorporated).

For detection of single mRNA molecules, we used the commercially available system RNAscope (ACDbio). Ntsr1-Cre;tdTomato mice (P23 to P30) were transcardially perfused with 0.1 M PBS followed by a 4% PFA solution, and brains were post-fixed for 2 h at room temperature in 4% PFA. Fixed brains were then washed in PBS and allowed to equilibrate in a 30% sucrose-PBS solution for 36 h before being embedded and frozen in OCT compound (Scigen, Tissue-Plus #4583). Frozen brains were kept at-80°C for up to 2 months. 20 μm sections were collected on a cryostat (Leica CM3050) and processed according to instructions provided by RNAscope. The following probes were used: tdTomato-C2 (317041-C2), tdTomato-C3 ( 317041-C3), Mm-Rbfox3 (313311), Rbfox3-C2 (313311-C2), Mm-Lamp5 (451071), Mm-Pantr1 (483711), Mm-Serpini1 (501441), Gabra5-C3 (319481-C3). Imaging was performed on a confocal microscope (LSM 510, Zeiss) using a 63x objective (NA 1.4). Sections from three animals were used for *Pantr1, Lamp5* and *Gabra5* and from four animals for *Serpini1.* For quantification of the number of mRNAs per neuron obtained using RNAscope, single plane images were first processed with the open-source software CellProfiler (Broad Institute) to automatically identify individual cells based on DAPI signal and to generate outline files. These outlines were then used as input, in combination with the corresponding confocal images, into FISH-quant^64^ (Matlab) for quantification of transcript number per cell. Cells with no detectable NeuN *(Rbfox3)* transcript were discounted as non-neuronal cells. A threshold of 20 *tdTomato* puncta was selected as the cutoff for distinguishing tdTomato-positive from tdTomato-negative neurons. This resulted in 164 tdTomato-positive and 188 tdTomato-negative neurons for *Serpini1,* 118 tdTomato-positive and 150 tdTomato-negative neurons for *Lamp5,* 111 tdTomato-positive and 198 tdTomato-negative neurons for *Pantr1,* and 135 tdTomato-positive and 160 tdTomato-negative neurons for *Gabra5.* Individual images were stitched to reconstitute an image spanning layer 6 vertically, and the distance of each neuron from the bottom of the layer was measured manually using Fiji^57^ and normalized to the height of layer 6. Statistical testing (Likelihood ratio test) and curve fitting (Loess) was performed in R/Bioconductor.

### Sensory manipulation

Prior to sensory manipulation, animals were bilaterally injected in POm with green Retrobeads IX (Lumafluor) as described above. To induce experience dependent plasticity, mice (P18 to P23) were anesthetized with the inhalation anesthetic, isoflurane. Once an adequate level of anesthesia was achieved, the animals were placed under a surgical microscope and, using tweezers, individual vibrissae were gently pulled by applying slow, steady tension to the base of the whisker^65^. To produce a chessboard pattern, vibrissae alpha, gamma, A2, A4, B1, B3, C2, C4, C6, D1, D3, D5, D7, E1, E3, E5 and E7 were pulled unilaterally while the rest remained untouched. Following the procedure, the animals were returned to their home cage and housed with at least one other animal until tissue collection. We checked for regrowing vibrissae every two days and removed them if visible.

### Read pre-processing

For both bulk and single-cell libraries, paired-end reads were aligned to the mouse reference genome (mm10) using Hisat2^66^ with the default parameters except:-p 8. Aligned reads from individual samples were quantified against a reference transcriptome^61^ (GENCODE vM8) supplemented with additional lncRNA genes described previously^8^ (Supplementary File S1) with cuffquant^67^ using default parameters with the following exceptions:‐‐no-update-check-p 8. For bulk RNA-Seq, pre-processed expression estimates were used as input for cuffdiff using default parameters except:-p 16, and the output of cuffdiff was analyzed using the cummeRbund R/Bioconductor package^68^. For single-cell experiments, normalized expression estimates across all samples were obtained using cuffnorm with default parameters. For comparison to results from primary visual cortex^12^, raw reads were downloaded from the short read archive (GSE71585), processed as described above for single-cell samples, and normalized expression estimates were obtained from combining both sources of single-cell reads together via cuffnorm.

### Single-cell analysis

The normalized FPKM matrix from cuffnorm was used as input for the Monocle2 single-cell RNA-seq framework^69^ in R/Bioconductor^70^. Relative FPKM values for each cell were converted to estimates of absolute mRNA counts per cell (RPC) using the Monocle2 census utility. A total of 346 individual cells under baseline conditions passed quality filters including: a) total mRNAs between 2000 and 200,000 per cell, and b) >10,000 fragments mapped (total_mass). 12,537 genes were identified as expressed in at least 15 cells across all conditions.

To identify high-variance genes, a generalized additive model^71^ (MGCV R package) was fit to the log_2_ mean RPC expression versus a cubic spline fit to the log_2_ coefficient of variation(BCV) for each replicate dissociation independently (Supplementary Fig. 2a and Fig. 7d). The intersection of genes with residuals to this fit greater than 1.0 from each replicate were chosen as ‘high-variance’ genes and the log_2_ expression estimates (with a pseudocount of 1) of these selected genes were used as input for PCA analysis and t-stochastic nearest neighbor embedding (tSNE) clustering of individual cells^72^.

To cluster individual cells, we employed a spectral tSNE approach^73^, combining nonlinear dimensional reduction and density clustering. Briefly, PCA analysis was conducted on high-variance genes and then a subset of components (n = 3, selected by permutation parallel analysis, Supplementary Fig. 2b) were used as input for the tSNE visualization algorithm^72^ (perplexity = 30). K-means clustering was used to partition cells using the two reduced tSNE dimensions into two discrete subtypes. The choice of k was made using 1000 bootstrapped tSNE estimates and evaluating k = 2 through k = 8 for each round. The average silhouette coefficient for each tSNE*k was determined and the mean and standard deviation of these estimates was estimated for each k (Supplementary Fig. 2c). k = 2 was chosen as the k value with the highest average silhouette coefficient and the lowest value of k for which there was no significant improvement in mean silhouette score.

After cluster assignment, differential expression testing was performed across all expressed genes using the Monocle2 VGAM model comparison test^69^. To test for differentially expressed genes between transcriptionally-defined cell types, the following full model was fit to each expressed gene: ~replicate+num_genes_expressed+celltype, and compared to a reduced model in which celltype was removed. The number of genes expressed in each cell was included as an explanatory variable as a proxy for the sensitivity of scRNA-seq in each individual cell. To test for differentially expressed genes between anatomically labeled cells, the following full model was fit to each expressed gene: ~replicate+num_genes_expressed+label, and compared to a reduced model in which label was removed. In both cases, genes were selected as significantly differentially expressed with a 0.1% false discovery rate (Monocle2 test; Benjamini-Hochberg corrected). DE gene lists were tested for gene set enrichment against Gene Ontology and Reactome genesets^74, 75^ using the clusterProfiler Bioconductor package^76^. The heatmap of differentially expressed genes was composed using the pheatmap R package^77^ and dendrograms were generated from the hierarchical clustering of the Euclidean distances between points.

To identify gene sets contributing to transcriptional identity in our single-cell dataset, gene-centric PCA was performed on a mean-centered matrix of variance-stabilized expression estimates for high-variance genes across all cells. The resulting rotations were used to project all expressed genes into the same PCA space to identify their weights. These weights were used to rank-order all expressed genes and this ordering was used as input for a pre-ranked GSEA analysis^78^. Gene sets for the GSEA analysis were derived from the Monocle2 differential gene tests described above or from our curated list of neuronal activity genes^43-46^ using their adjusted p-value cutoff of p < 0.01 (Supplementary Table 4).

### WGCNA

To identify modules of correlated gene expression we used the Weighted Gene Correlation Network Analysis package WGCNA^42^. Normalized expression data were first batch-corrected with respect to replicates using limma, and the resulting matrix of 12,537 detectably expressed genes and 346 cells was used as input for WGCNA. Cells were hierarchically clustered using the Jensen-Shannon distance with average linkage. A soft threshold power of 11 was used to create a signed Topological overlap matrix directly from the normalized and batch-corrected expression estimates. Modules were then learned with a minimum module size of 30. Modules were merged below a distance threshold of 0.5. Module eigenvalues were estimated and correlated with cellular traits using the Pearson product moment correlation test. Module gene membership was determined in a similar manner in which individual gene expression estimates were correlated with module eigenvalues and tested for significance. To test the effect of each module on the segregation of L6CThN cell types, each module eigenvalue was separately regressed out of the expression matrix using limma, and subsequent values were used as input for a tSNE using identical parameters to the original assay.

### Pseudotime and BEAM analyses

Pseudotemporal ordering was performed using the prescribed Monocle2 workflow. Briefly, all 1023 L6CThNs passing quality controls were used as input. To reconstruct a trajectory that reflected cellular progression in response to altered sensory input, we first performed a differential test to identify genes whose expression changed as a function of collection day, independent of baseline differences between celltype (fullModelFormulaStr = "~num_genes_expressed+Total_mRNAs+sex+celltype*Day, reducedModelFormulaStr = " ~num_genes_expressed+Total_mRNAs+sex+celltype", q ≤ 0.01). These 1134 genes were used as a filtering set for the DDRTree dimensionality reduction. Cells were tested for differential expression along pseudotime using the following model comparisons with q ≤ 0.0000001: fullModelFormulaStr = " ~num_genes_expressed+celltype+sm.ns(Pseudotime,df=3)", reducedModelFormulaStr = " ~num_genes_expressed+celltype". Branch dependent expression was determined using the Monocle2 BEAM test (fullModelFormulaStr = " ~num_genes_expressed+celltype+sm.ns(Pseudotime, df = 3)*Branch", reducedModelFormulaStr =" ~num_genes_expressed+celltype+sm.ns(Pseudotime, df = 3)", q ≤ 0.0001). All significant gene lists were tested for gene set enrichment using the hypergeometric test.

### Data and software availability

All primary data are archived on the SRA and Gene Expression Omnibus and are available for direct download on our companion website (http://rstudio.gofflab.org:8383/apps/L6CthPN_dash/). Source code and software tools are available upon request.

## Author Contributions

MC, SPB and LAG conceived the experiments. MC, JDR, and GHC performed all experiments. MC, SPB and LAG analyzed the data and wrote the paper with input from all authors.

## Conflict of Interest

The authors declare no competing financial interests.

## Acknowledgments

We thank Hao Zhang and the Johns Hopkins Bloomberg School of Public Health Flow Cytometry and Cell Sorting Core Facility. Sequencing service was provided by the Johns Hopkins Genetic Resources Core Facility. We thank Dr. Joshua R. Sanes, Harvard University, for the anti-LacZ antibody and Dr. Jesse Gray, Harvard Medical School, for the curated list of neuronal activity-induced genes. This work was supported by a Klingenstein-Simons Fellowship (SPB), a Johns Hopkins Science of Learning Grant (SPB, LAG), National Science Foundation Grant 1656592 (SPB, LAG), NIH training grant T32 GM07814 (JDR), a Boerhinger-Ingelheim Fonds Fellowship (MC), and the NINDS (NS050274).

